# Probabilistic Framework for Integration of Mass Spectrum and Retention Time Information in Small Molecule Identification

**DOI:** 10.1101/2020.08.19.255653

**Authors:** Eric Bach, Simon Rogers, John Williamson, Juho Rousu

## Abstract

**Motivation:** Identification of small molecules in a biological sample remains a major bottleneck in molecular biology, despite a decade of rapid development of computational approaches for predicting molecular structures using mass spectrometry (MS) data. Recently, there has been increasing interest in utilizing other information sources, such as liquid chromatography (LC) retention time (RT), to improve the MS based identifications.

**Results:** We put forward a probabilistic modelling framework to integrate MS and RT data of multiple features in an LC-MS experiment. We model the MS measurements and all pairwise retention order information as a Markov random field and use efficient approximate inference for scoring and ranking potential molecular structures. Our experiments show improved identification accuracy by combining tandem mass spectrometry data (MS^2^) and retention orders using our approach, thereby outperforming state-of-the-art methods. Furthermore, we demonstrate the benefit of our model when only a subset of LC-MS features have MS^2^ measurements available besides MS^1^.

**Availability and implementation:** Software and data is freely available at https://github.com/aalto-ics-kepaco/msms_rt_score_integration.

## 1 Introduction

The identification of small molecules, such as metabolites or drugs, in biological samples is a challenging task posing a bottleneck in various research fields such as biomedicine, biotechnology, environmental chemistry and drug discovery. In untargeted metabolomics studies, the samples typically contain thousands of different molecular species, the vast majority of which remain unidentified (Aksenov *et al*., 2017; da Silva *et al*., 2015). Liquid chromatography (LC) coupled with tandem mass spectrometry (MS^2^) is arguably the most important measurement platform in metabolomics (Blaženovi ć *et al*., 2018), due to its suitability to high-throughput screening, its high sensitivity and applicability to a wide range of molecules. Briefly explained, LC separates molecules by their retention time (RT) and MS separates molecules by their mass (MS^1^). Subsequently, MS^2^ can be used to fragment molecules in a narrow mass window and to record the fragment intensities (MS^2^-spectrum). In an untargeted metabolomics experiment, large sets of MS features (MS^1^and RT, plus optionally MS^2^), are observed, corresponding to the different molecular species in the sample. Metabolite identification concerns then the structural annotation of the observed MS features.

In recent years numerous powerful approaches (Nguyen *et al*., 2018a; Schymanski *et al*., 2017) for annotating MS^2^ spectra with a predicted molecular structure have been developed (Ruttkies *et al*., 2016, 2019; Dührkop *et al*., 2015; Brouard *et al*., 2016; Allen *et al*., 2014; Nguyen *et al*., 2018b, 2019; Dührkop *et al*., 2019). Typically, these methods output a ranked list of molecular structure candidates, that can be shown to human experts, or further post-processed, e.g. by using *additional* information available for the analysed sample. Sources of additional information include, e.g. RT (Ruttkies *et al*., 2016; Bach *et al*., 2018; Samaraweera *et al*., 2018), collision cross-section (Plante *et al*., 2019), or prior knowledge on the data generating process, such as the source organism’s metabolic characteristics (Rutz *et al*., 2019).

Retention time, that is, the time that a molecule takes to elute from the LC column, is readily available in all LC-MS pipelines, and is frequently used in aiding annotation (Stanstrup *et al*., 2015). A basic technique is to use the difference between the observed and predicted RT (Samaraweera *et al*., 2018; Domingo-Almenara *et al*., 2019) to prune the list of canididate molecular structures. A major challenge for utilizing RT information, however, is that the RT of the same molecule can vary significantly across different LC systems and configurations, necessitating system specific candidate RT reference databases and RT predictors. Different approaches have been proposed to tackle this challenge, such as using physico-chemical properties (e.g., partition coefficient, LogP) as RT proxies (Ruttkies *et al*., 2016; Hu *et al*., 2018), retention time mapping across LC systems (Stanstrup *et al*., 2015) or predicting retention orders, which are largely preserved within a family of LC systems (e.g. reversed phase) (Bach *et al*., 2018; Liu *et al*., 2019). Using LogP as an RT proxy is simple to implement, but only models the hydrophobic separation effects of the LC system. RT mapping is limited to LC system pairs, restricting the number of available training samples (i.e. the subset of structures observed in both systems). Retention order prediction can overcome those drawbacks, by learning the LC system’s separation directly from RT data of multiple systems (Bach *et al*., 2018).

This study proposes a probabilistic framework to integrate MS^1^ or MS^2^ based annotations with predicted retention order for improved small molecule identification given a set of MS features by building on the work by Bach *et al.* (2018) and Del Carratore *et al.* (2019). The latter proposed a *probabilistic* approach for integrating different types of additional information to MS^1^ data, including RT information. We too define a probabilistic approach, but differ in how RT is handled. Where Del Carratore *et al.* (2019) use absolute RT information, we follow Bach *et al.* (2018) and use pairwise retention order predictions. In contrast to the work done by Bach *et al*. (2018), our model makes use of pairwise retention order information between all candidate molecular structures, resulting in more accurate annotations and allowing us to re-rank all candidates lists, rather than just returning the most likely candidate assignments for each MS feature.

Our framework models the score integration as an inference problem on a graphical model, where the edges correspond to retention order predictions, the nodes correspond to MS features, and the node labels correspond to candidate molecular structures, scored by a MS^2^ based predictor, such as CSI:FingerID (Dührkop *et al*., 2015), MetFrag (Ruttkies *et al*., 2016) or IOKR (Brouard *et al*., 2016), or in the absence of MS^2^ information, MS^1^ precursor mass deviation. This graph is fully connected, which makes exact inference an NP-hard problem. To solve this challenge, we resort to efficient approximate inference, in particular spanning tree approximations (Wainwright *et al*., 2005; Pletscher *et al*., 2009; Su and Rousu, 2015; Marchand *et al*., 2014).

## 2 Methods

### 2.1 Overall workflow

We assume data arising from a typical LC-MS based experimental workflow (including chromatographic peak picking, and alignment): MS features consisting MS^1^ measurement and the associated retention time (RT). A subset of these will include an MS^2^ spectrum. In the following, we present our score integration model in the most general form in which it is provided with MS features and a set of possible candidate molecular structures. The candidate list can be generated, for example, by querying molecular structures from a structure database such as ChemSpider (Pence and Williams, 2010) that have the same mass as the observed MS feature. In addition, we assume that to each candidate structure a score is assigned by either an MS^2^ based predictor, or, if no MS^2^-spectrum is available, a score based on the mass deviation of the candidates from the MS mass. For all molecular candidate pairs, associated with the different MS features, the retention order is predicted. Here we use the Ranking Support Vector Machine (RankSVM) based predictor by Bach *et al.* (2018). The candidate structure scores and predicted retention orders are integrated through a probabilistic graphical model (described in the following). This allows us to rank the molecular candidate structures by their inferred marginal probabilities, given both the MS and RT information.

More formally, the output of an LC-MS experiment is given as a tuple set 𝒟 = *{*(*x*_*i*_, *t*_*i*_, 𝒞 _*i*_) *}*, with *x*_*i*_ ∈ 𝒳 being the spectrum of feature *i* (either an MS^2^ or a spectrum containing only a single peak at the mass of the precursor ion if no MS^2^ information is available), *t*_*i*_ ∈ℝ_≥0_ being its retention time, and 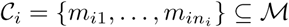 being the associated molecular candidates. Here, *m*_*ir*_ ∈ ℳ represents a molecular candidate structure, and *n*_*i*_ is the number of molecular candidates for the *i*th MS feature. Figure 1 shows an overview of our workflow.

**Figure 1:**
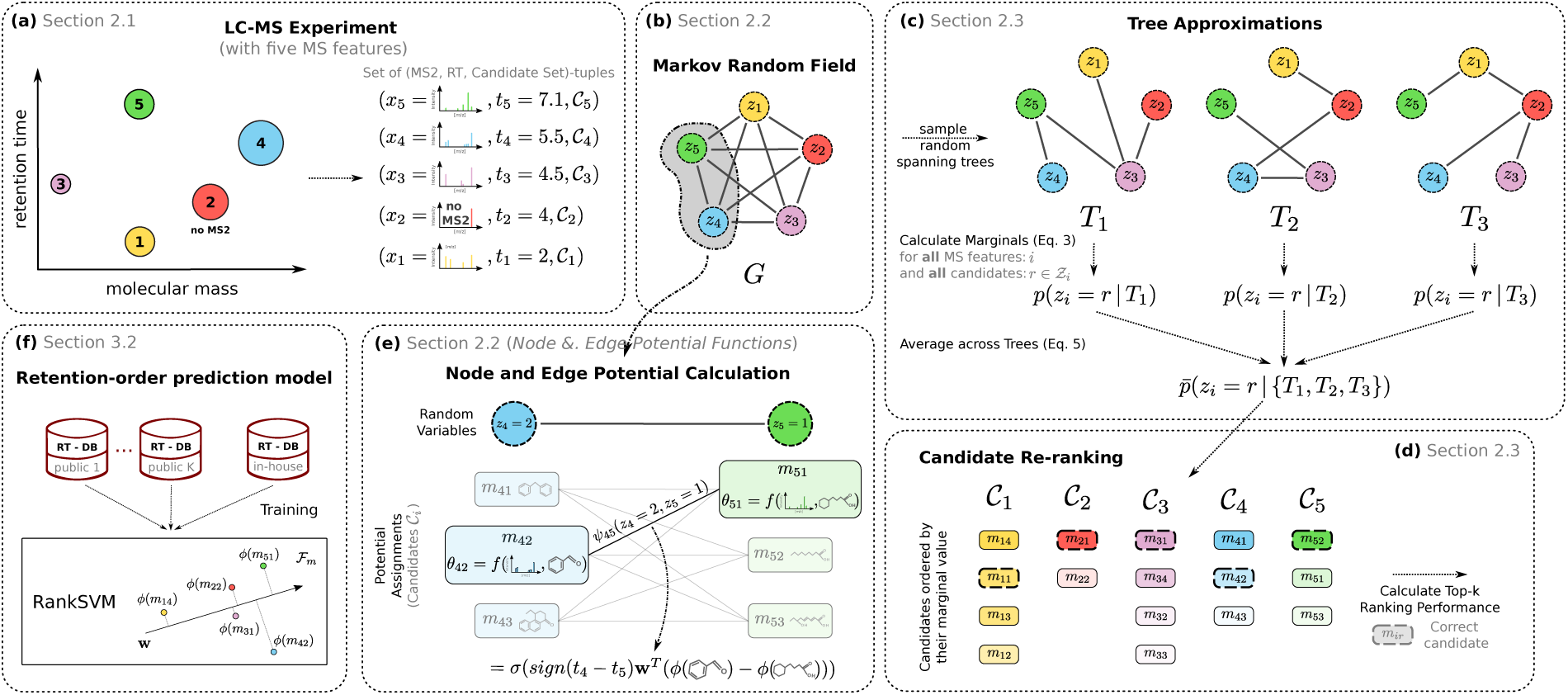
Workflow of our framework and its main components. (a) Data acquisition in an LC-MS experiment resulting in a set of (MS^2^, RT)-tuples of unknown molecules. (b) Illustration of the underlying graphical model. (c) Ensemble of spanning trees to approximate the markov random field and their integration using averaged marginals. (d) Output to the user: Re-ranked molecular candidate lists based on the approximated marginals. (e) Incorporation of the predicted retention orders for a particular assignment for **z** via the edge potential function. (f) Illustration of the RankSVM model.

### 2.2 Probabilistic Model

Let *G* = (*V, E*) be an undirected graph, in which each node, *i* ∈*V* represents one observed MS feature, and with an edge for all MS feature pairs *E* = *{* (*i, j*) | *i, j* ∈ *V, i* ≠ *j}*. The edge set *E* does not contain any parallel edges. The number of MS features is denoted with *N*, i.e. |*V* | = *N*. We associate each node *i* in the vertex set with a discrete random variable *z*_*i*_ that takes values from the space 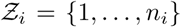. Intuitively, *z*_*i*_ defines which candidate has been assigned to the *i*th MS feature. The full vector **z** = *{z*_*i*_ |*i* ∈ *V}* corresponds to the molecular structure assignment to each MS feature in the LC-MS experiment, and it takes values from the set 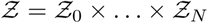. In this work we consider 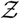 to be fixed and finite for a given a set of MS features, due to our definition of the molecular candidates sets which assumes that we can restrict the putative annotation for a given MS feature.

#### Markov Random Field

The probability distribution of **z** is given as a *pairwise Markov Random Field* (MRF) (MacKay, 2005):

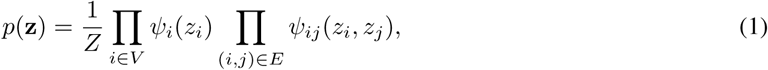

composed of node *ψ*_*i*_ and edge *ψ*_*ij*_ potential functions, and omits higher-order cliques (hence the term pairwise). Above, 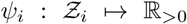 is the potential function of node *i* measuring how well the *i*’s candidates matches the measured MS information, and 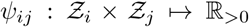 encodes the consistency of the observed retention orders for MS feature *i* and *j* and the *predicted* retention order of their candidates *z*_*i*_ and *z*_*j*_ and 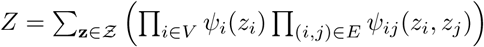 is the partition function (MacKay, 2005).

#### Node Potential Function *ψ*_*i*_

For each candidate 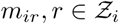 we predict a matching score *θ*_*ir*_ = *f* (*x*_*i*_, *m*_*ir*_) ∈ ℝ expressing how well it matches the observed MS^1^ or MS^2^ spectrum *x*_*i*_. For that, we assume a pre-trained model, such as CSI:FingerID (Dührkop *et al*., 2015), MetFrag (Ruttkies *et al*., 2016) or IOKR (Brouard *et al*., 2016). We use the latter two in our experiments as representative MS^2^ scoring methods (Sec. 3.3). MetFrag performs an in-silico fragmentation of *m*_*ir*_, compares these fragments peaks with the observed ones in *x*_*i*_ and outputs a matching score. IOKR, on the other hand, can be used to directly predict a matching score *f* (*x, m*) for any (MS^2^ feature, molecular structure)-tuple. All matching scores *θ*_*ir*_ are normalized to the range [0, 1]. Finally, we express the potential of a molecular candidate *m*_*r*_ given the spectrum *x*_*i*_ as follows:

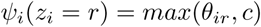

where *c* > 0 is a constant used to avoid zero potentials. In our experiments, we select *c* such that it is ten times smaller than the minimum of all non-zero scores across all candidate sets.

#### Edge Potential Function *ψ*_*ij*_

For each candidate pair 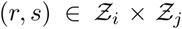 associated with the MS pair (*i, j*) we compute how well the candidates’ predicted retention order is aligned with the observed one defined by the retention times *t*_*i*_ and *t*_*j*_. To this end, we apply the framework for retention order prediction developed by Bach *et al.* (2018). The edge potential *ψ*_*ij*_(*z*_*i*_ = *r, z*_*j*_ = *s*) is defined as follows:

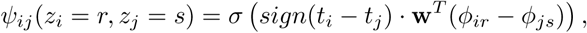

where 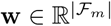 is the RankSVM’s parameter vector, and *ϕ*_*ir*_, *ϕ*_*js*_ ∈ ℱ _*m*_ are the feature vectors of the candidates’ molecular structures, and *σ*: ℝ ↦ (0, 1] is a monotonic function mapping the predicted preference value difference to a value between zero and one. In our experiments we consider two mapping functions:

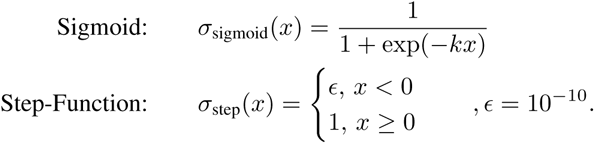

The different functions can be interpreted as follows. The *sigmoid* makes full use of the information from the RankSVM margin, i.e. the score of each candidate pair depends on the preference score difference. In this work, we consider *k* as a hyper-parameter of our method, that needs to be estimated from data (Sec. 3.5). The *step-function*, on the other hand, only differentiates between aligned and not aligned pairs.

#### Weighting of Information Sources

To control the contribution of each information source, i.e. MS information and retention orders, we introduce a modification on the potential functions:

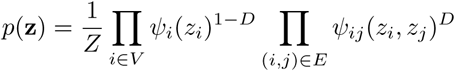

with *D* ∈ [0, 1]. A *D* value close to one, for example, will result in a score mainly based on the observed retention orders. In our experiments we explain how this hyper-parameter can be estimated in practice (Sec. 3.5).

### 2.3 Ranking Candidates through approximated Marginals

We rank the molecular candidates using the marginals of the MRF (1). The marginal for the candidate *r* of MS feature *i* is given as:

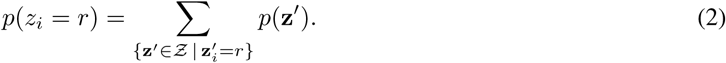

In practice, the calculation of (2) is intractable due to the size of the domain 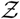 of **z**, which grows exponentially with the number of MS features, thus we will resort to approximate inference methods.

#### Tree Approximation of *G*

To enable feasible inference of (2) we approximate the MRF (1) using spanning trees of the original graphical model *G* (Wainwright *et al*., 2005; Pletscher *et al*., 2009; Su and Rousu, 2015; Marchand *et al*., 2014). In the following let *T* be a spanning tree of *G* with the same nodes, but an edge-set *E*(*T*) *⊆E*, with |*E*(*T*) | = *N* − 1, that ensures *T* being a cycle-free single connected component. The probability distribution of the graphical model induced by *T* is given as:

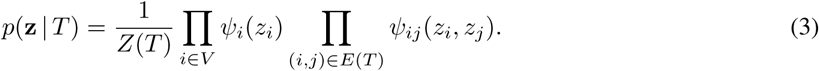

As the graphical model associated with (3) is a tree, we can exactly infer its marginals through the *sum-product* algorithm (MacKay, 2005), The sum-product algorithm is a message-passing algorithm using dynamic programming that has linear time complexity in the number of MS features. See, for example, MacKay (2005) for further details on the algorithm.

The output of the sum-product algorithm are the unnormalized marginals *µ*(*z*_*i*_ = *r*|*T*_*t*_) for all *i* ∈ *V* and 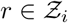. We calculate the normalized marginals as follows (MacKay, 2005):

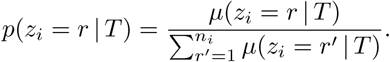

#### Random Spanning Trees Sampling

We compare two approaches to retrieve spanning trees from *G*. The first approach is to randomly sample spanning trees from *G* (c.f. Pletscher *et al*., 2009; Wainwright *et al*., 2005; Su and Rousu, 2015). We sample the trees by applying the minimum weighted spanning tree algorithm to a random adjacency matrix. If for an MS feature pair (*i, j*) both retention times are equal, i.e. *t*_*i*_ = *t*_*j*_, than their corresponding edge is not sampled. This is justified by the observation, that MS features with a retention time difference equal zero, do not impose constraints on the retention order of their corresponding candidates. We will refer to a sampled spanning tree as *T*_*t*_. The second approach was implicitly used by Bach *et al.* (2018) and corresponds to a linear Markov chain where edges connect adjacent MS features ordered by increasing RT, which can be seen as a degenerate spanning tree. In the remaining text we refer to this tree as *T*_chain_.

#### Averaged Marginal over a Random Spanning Tree Ensemble

Using tree-like graphical models for the inference is motivated by the exact and fast inference it enables us to do. However, a single tree, such as *T*_chain_ or a sampled *T*_*t*_, will most likely be only a rough approximation of the original probability distribution (1). Therefore, the use of random spanning tree ensembles 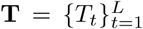 has been proposed. In particular, Wainwright *et al*. (2005) show that an expectation over trees can be used to obtain an upper bound on the maximum a posteriori (MAP) estimate of the original graph, and showed that this approximation can be tight if the underlying trees agree about the MAP configuration. More recently, Marchand *et al.* (2014) demonstrated generalization bounds for joint learning and inference using tree ensembles. More applied work in using tree-based approximation can be found in (Pletscher *et al*., 2009) who use majority voting and (Su and Rousu, 2015) who empirically study several aggregation schemes in multilabel classification.

Motivated by the mentioned literature, we opted to average the marginals of an random spanning tree ensemble **T**, where for each tree *T*_*t*_ we independently retrieve the marginals using the sum-product:

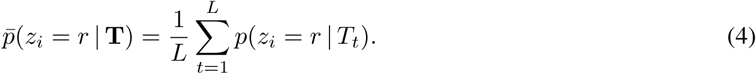

#### Max-Marginals

The exact inference on trees allows us to use the max-marginal, as an alternative to the sum-marginal shown in Equation (2). The max-marginal is closely related to the MAP estimate. For a single tree *T* it is given as:

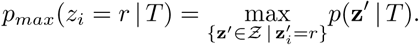

The interpretation of the two marginals (sum and max) differs slightly. Whereas the sum-marginal expresses the *sum* of the probabilities of all configurations 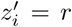, the max-marginal is the maximum probability that a configuration with the constraint 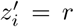 can reach. In our experiments we compare the performance of both marginal types (Sec. 4.1). The *max-product* algorithm is used to calculate the unnormalized max-marginals *µ*_*max*_(*z*_*i*_ = *r* | *T*), which is a modification of the sum-product algorithm, in which summations are replaced by maximization. The normalized marginal can be calculated as (MacKay, 2005):

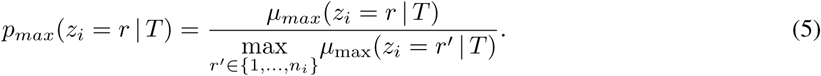

By plugging Equation (5) into (4) instead of the sum-marginal we obtain the averaged max-marginal 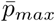.

#### Run-time Complexity

Calculating the marginals for an individual tree and all MS features *i* has run-time complexity 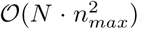. Here, *N* is the total number of features and 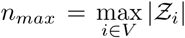 the maximum number of molecular candidates for a feature.

## 3 Material and Experiments

### 3.1 Evaluation datasets

To evaluate our score integration approach, we use two publicly available datasets. These are described in this section and summarized in Table 1.

**Table 1:** Summary of the datasets used for the evaluation of our score integration framework.

| Dataset | Ion. | Mass spectra info. |  | Molecular candidates<sup>a</sup> |  | Chromatography |  |
| --- | --- | --- | --- | --- | --- | --- | --- |
|  |  | MS<sup>1</sup> info. | #MS<sup>2</sup> | Tot. #Cand. | Med. #Cand. | Column | Eluent<sup>b</sup> |
| CASMI 2016 | neg. | Precursor m/z | 81 | 74589 | 420 | Phenomenex Kinetex EVO C18 | H2O → MeOH |
| CASMI 2016 | pos. | Precursor m/z | 127 | 183633 | 919 | Phenomenex Kinetex EVO C18 | H2O → MeOH |
| EA (Massbank) | neg. | Precursor m/z | 154 | 75107 | 119.5 | Waters XBridge C18 | H2O → MeOH |
| EA (Massbank) | pos. | Precursor m/z | 319 | 215893 | 246 | Waters XBridge C18 | H2O → MeOH |

<sup>a</sup> ChemSpider. CASMI: $\pm 5$ ppm around monoisotopic exact mass of correct candidate. EA: MF of correct candidate.
<sup>b</sup> Additive: Both 0.1% formic acid

#### CASMI 2016

The Critical Assessment of Small Molecule Identification (CASMI) challenge is a contest organized for the computational spectrometry community (Schymanski *et al*., 2017). For its implementation in 2016, a dataset of 208 (MS^2^, retention-time)-tuples was released. The dataset contains 81 negative and 127 positive ionization mode MS^2^ spectra. The molecular candidate structure sets were extracted from ChemSpider, using a *±*5ppm window around the monoisotopic exact mass of the correct candidate, by the challenge organizers.

#### EA (Massbank)

Massbank (Horai *et al*., 2010) is a publicly available repository for mass spectrometry data. For the development of MetFrag 2.2, Ruttkies *et al.* (2016) extracted 473 (MS^2^, retention-time)-tuples of 359 unique molecular structures from Massbank (EA dataset). The dataset is split into 154 negative and 319 positive ionization mode MS^2^ spectra. We used the molecular candidates provided by Ruttkies *et al.* (2016) extracted from ChemSpider using the molecular formula of the correct candidate.

For each dataset and ionization mode we repeatedly subsample training and test (MS^2^, RT)-tuple sets: CASMI (negative) 50-times *N*_*train*_ = 31, *N*_*test*_ = 50; CASMI (positive) 50-times *N*_*train*_ = 52, *N*_*test*_ = 75; EA (negative) 50-times *N*_*train*_ = 45, *N*_*test*_ = 65; and EA (positive) 100-times *N*_*train*_ = 50, *N*_*test*_ = 100. No molecular structure, determined by its InChI representation, appears simultaneously in test and training. The training set is used for the hyper-parameter selection (Sec. 3.5) and the test sets are used to assess the average identification performance of our score integration framework (Sec. 3.4).

### 3.2 Training Setup for the Retention Order Predictor

To calculate the edge potentials of our MRF model (1) we use the RankSVM retention order prediction approach by Bach *et al*. (2018). The RankSVM model is trained using seven publicly available retention time datasets. Six where published by Stanstrup *et al.* (2015) along with their retention time mapping tool PredRet: `UFZ_Phenomenex`, `FEM_long`, `FEM_orbitrap_plasma`, `FEM_orbitrap_urine`, `FEM_short` and `Eawag_XBridgeC18`. The seventh dataset contains examples for which retention times were published as part of the training dataset for the CASMI 2016 challenge (Schymanski *et al*., 2017). The joint dataset covered four different chromatographic columns all using H2O *→* MeOH (with 0.1% formic acid as additive) as eluent. In total, the dataset contained 1248 (molecule, RT)-tuples of 890 unique molecular structures, after the same pre-processing as in Bach *et al.* (2018) was applied. We represent the molecular structures using `Substructure` counting fingerprints calculated with `rcdk` and `CDK` 2.2 (Willighagen *et al*., 2017). We use the MinMax-kernel (Ralaivola *et al*., 2005) to calculate the similarity between the fingerprints. For our experiments we build an individual RankSVM model for each (MS, RT)-tuple subsample (Sec. 3.1), ensuring no molecular structure in the subsample is used for the RankSVM training.

### 3.3 MS^2^-based match scores from MetFrag and IOKR

We apply MetFrag (Ruttkies *et al*., 2016) and IOKR (Brouard *et al*., 2016) as representative methods to obtain MS^2^ matching scores for the molecular structures in the candidate list of each MS^2^ spectrum.

#### MetFrag

We use the latest MetFrag version 2.4.5 ^1^ and utilize it as described in Ruttkies *et al.* (2016). The MS^2^ matching scores are calculated using the `FragmenterScore` feature of MetFrag.

#### IOKR

Two IOKR models are trained, for negative and positive mode MS^2^ spectra, respectively. The training (MS^2^, molecular structure)-tuples are extracted from GNPS (Wang *et al*., 2016), Massbank and the CASMI 2016 training data. We remove training molecular structures that appear in our evaluation datasets (Sec. 3.1). This results in 3255 negative and 6773 positive mode training examples. We use a uniform combination of 16 MS^2^ spectra and fragmentation tree (FT) kernels as input kernel (supplementary material, Sec. S.4). On the output side, we use the same molecular fingerprint definitions as Dührkop *et al.* (2019) as feature representation and a Gaussian kernel those distances are derived from the Tanimoto kernel (Brouard *et al*., 2019) as output kernel. For all MS^2^ spectra in our evaluation datasets, we calculate the FTs using SIRIUS 4.0.1 (Dührkop *et al*., 2019) and keep the highest scoring tree for each spectra to calculate the MS^2^ and FT kernels used by the IOKR.

### 3.4 Performance Evaluation

In our experiments, we use the top-*k* accuracy to determine the metabolite identification performance, that is, the percentage of correctly ranked molecular candidates at rank *k* or less. Different approaches can be used to determine the rank of the correct structure. We follow the protocol used by Schymanski *et al.* (2017). If multiple stereo-isomers were present in the candidate list, only the one with the highest MS^2^-score was retained. The correct molecular structure was found by comparing the InChIs containing no stereo information. The top-*k* accuracies are calculated the test sets.

### 3.5 Hyper-parameter Estimation

The training set of each individual subsample is used to determine optimal weighting *D* between MS and retention order information. For that, we run the score integration framework for a different *D* values, and calculate the area under-the-ranking curve up to rank 20: 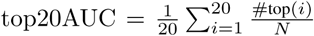, where #top(*i*) is the number of correct structures up to rank *i*, and *N* is the number of MS features. Subsequently we select the retention order weight with the highest top20AUC (supplementary material, Sec. S.2). The optimal sigmoid parameter *k* is estimated using Platt’s method (Platt, 2000; Lin *et al*., 2007) calibrated using RankSVM’s training data (Sec. 3.2).

### 3.6 Experiments

#### Full MS^2^ Information available

We compare our approach for combining MS^2^ and RT information for metabolite identification against the baseline, which only uses MS^2^information for the candidate ranking. This allowed us to quantify the performance gain by using RTs. Furthermore, we applied two recently published methods for the integration of MS^2^ and RT scores and compared them to our approach. The first one is MetFrag 2.2 (Ruttkies *et al*., 2016) which exploits the RT information by establishing a linear relationship between the candidates’ predicted LogP values with the observed retention times. Each molecular candidate receives an additional score by comparing its predicted retention time against that observed for the corresponding MS^2^ spectra. We use the CDK (Willighagen *et al*., 2017) XLogP predictions which are automatically calculated by the MetFrag software. The weight of the retention time feature `RetentionTimeScore` is determined as described in Section 3.5. Our second comparison is the approach by Bach *et al.* (2018), which uses predicted retention order and dynamic programming over a chain graph connecting adjacent MS features. Bach *et al.* (2018) focused in extracting the most likely assignment **z** (Eq. (3)) given the chain graph using dynamic programming. Here, we use our generalized framework to also compute the marginals of all candidates given the chain graph *T*_*chain*_. The approach by Bach *et al.* (2018) implicitly used a hinge-sigmoid to compute edge potentials: 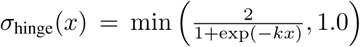. Its parameter *k* is determined as described in the supplementary material (Sec. S.2). We refer to this method as Chain-graph.

#### Missing MS^2^

In a second experiment, we simulated the common application scenario, in which during an LC-MS experiment a set of MS features (MS and RT) have been measured, but MS^2^ spectra have only been acquired for a subset of the features. There can be multiple reasons for this, such as limited measuring time when using, for example Data-Dependent Aquisition (DDA, Xiao *et al.* (2012)) protocols, bad fragmentation quality, or inability to deconvolute all spectra when using Data-Independent acquisition (DIA)). In this case, besides the RT, only MS^1^ related information is available for some features, which includes the mass of the ion (precursor m/z) and its isotope pattern. We use our proposed score-integration framework to perform structural identification when the proportion of MS features that have an MS^2^spectrum varies. We vary the percentage of available MS^2^-spectra from 0% to 100% and investigate how the joint use of MS^1^, MS^2^ and RT information can improve over the baseline solely relying on only MS information. For the candidates *r* of an MS feature *i*, simulated to be without MS^2^spectrum, we assign the following candidate score (Del Carratore *et al*., 2019):

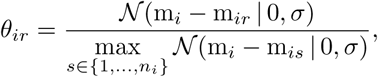

where m_*i*_ is the precursor mass of the measured ion, and m_*ir*_ is the mono-isotopic mass of candidate *r* associated with MS feature *i*, and 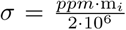 variance of Gaussian noise model. *ppm* expresses the MS-device accuracy, which we set to *ppm* = 5.

## 4 Results

### 4.1 Parameters of our Framework

This section investigates the influence of different settings for framework, such as number of random spanning trees or the marginal type.

#### Number of Random Spanning-Trees and Marginal Type

Figure 2 shows the top-*k* accuracy as a function of the number of random spanning-trees *L* averaged across the datasets and ionizations. The identification performance increases for larger *L*, whereby the improvement per tree decreases. For the top-1 performance remains similar for *L* ≥ 16 trees. However, for top-20 we observe improvements till *L* = 128. Figure 2 also shows that the max-marginal approach performs slightly better than the sum-margin. An explanation could be that max-marginal is more robust against candidate configurations **z** with very low probability. The sum-marginal averages over such cases, whereas the max-marginal only includes the one with maximum probability.

**Figure 2:**
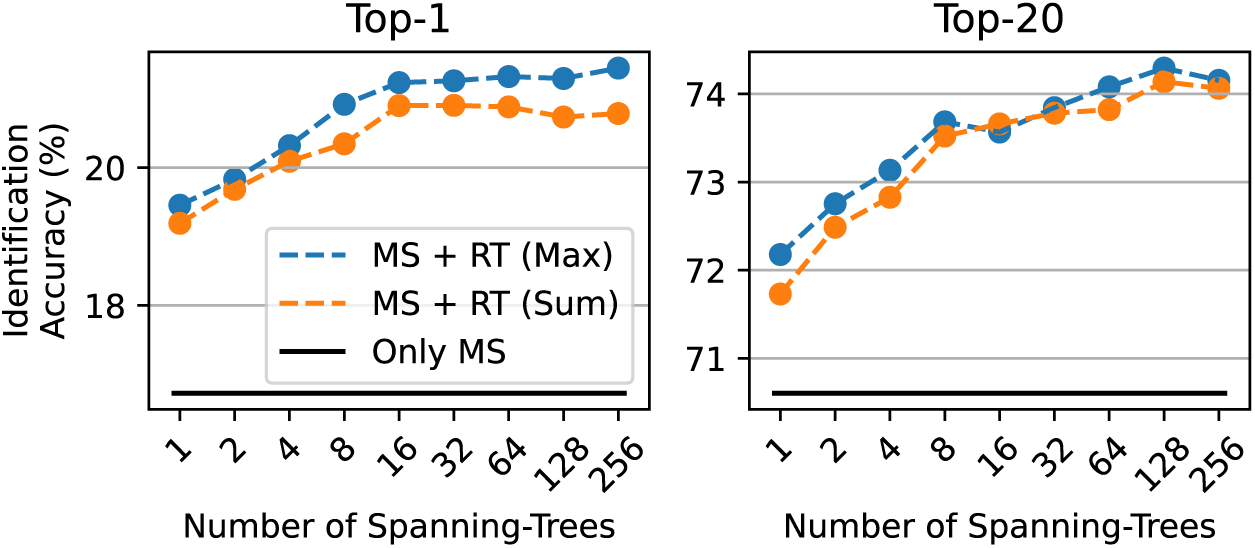
Top-*k* accuracies, averaged across all datasets and ionizations, plotted against the number of random spanning-trees used for the approximation. The baseline using only MS^2^ information is plotted in black. The sigmoid function is used in the score-integration.

#### Comparison of the Edge Potential Functions

The average metabolite identification performance does not differ much between two edge potential functions (see Table S1 in supplementary material). This is interesting specifically for the Step-function, which uses the predicted retention orders in a binary fashion only. However, the Sigmoid function still can significantly outperform the Step-function for top-1 and top-5 accuracy.

### 4.2 Performance of our Score Integration Framework

Here we compare our score integration framework with other methods and evaluate it under different data setups. we use *L* = 128 with max-marginals and the Sigmoid as edge potential function for the experiments.

#### Comparison to other Approaches

In Table 2 we compare the performance of our score integration framework with other approaches from the literature that utilize RT information for metabolite identification. It can be seen that our framework performs well across all datasets and ionization modes and we reach significant improvements over the baseline (Only MS). Especially for the positive mode spectra, our method seems to have an advantage, as both competing approaches, cannot consistently improve the identification by including RT information. The least performance improvement of our approach can be observed for the negative CASMI dataset, which might be due to the small training set. The other approaches, MetFrag 2.2 and Chain-graph, can consistently (top-1, 5, 10 and 20) improve the results only on particular (dataset, ionization mode) combinations. However, they almost always increase top-1 performance. The results in Table 3 show that our framework significantly outperforms MetFrag 2.2 and Chain-graph in terms of identification performance.

**Table 2:** Identification accuracies (top-*k*) for the different datasets and ionization modes. Compares our score integration framework (MS + RT (our)), against the baseline (Only MS), MetFrag 2.2 with predicted RT and the Chain-graph model. The best performance for each dataset and ionization is indicated by bold-font. The stars (*) represent the significant ^a^ improvement over the baseline calculated using a one-sided Wilcoxon signed-rank test on the sample top-*k* accuracies.

| Ionization | Dataset | Method | Top-1 | Top-5 | Top-10 | Top-20 |
| --- | --- | --- | --- | --- | --- | --- |
| Negative | CASMI 2016 | MS + RT (*our*) | **15.2** (***) | 47.2 (***) | 57.0 (**) | 70.1 (***) |
|  |  | MS + RT (Chain-graph) | 13.2 (***) | **49.4** (***) | **61.0** (***) | 69.4 (***) |
|  |  | MS + RT (MetFrag 2.2) | 14.0 (***) | 42.0 | 55.5 | **71.2** (***) |
|  |  | Only MS | 11.1 | 44.2 | 55.3 | 68.0 |
|  | EA Massbank | MS + RT (*our*) | 28.7 (***) | **61.9** (***) | **73.8** (***) | 83.6 (***) |
|  |  | MS + RT (Chain-graph) | 27.2 (***) | 59.5 (***) | 72.4 (***) | 81.8 (***) |
|  |  | MS + RT (MetFrag 2.2) | **30.2** (***) | 59.2 (***) | 73.6 (***) | **84.4** (***) |
|  |  | Only MS | 22.8 | 57.6 | 69.5 | 78.5 |
| Positive | CASMI 2016 | MS + RT (*our*) | **14.0** (***) | **40.7** (***) | **52.2** (***) | **62.8** (***) |
|  |  | MS + RT (Chain-graph) | 11.9 | 36.5 | 50.2 (***) | 60.7 (***) |
|  |  | MS + RT (MetFrag 2.2) | 13.7 (***) | 36.2 | 46.2 | 57.5 |
|  |  | Only MS | 11.8 | 37.3 | 47.0 | 58.3 |
|  | EA Massbank | MS + RT (*our*) | **27.3** (***) | **61.6** (***) | **72.9** (***) | **80.7** (***) |
|  |  | MS + RT (Chain-graph) | 23.9 (***) | 59.2 | 70.1 | 79.1 (***) |
|  |  | MS + RT (MetFrag 2.2) | 24.0 (***) | 59.0 | 69.5 | 77.1 |
|  |  | Only MS | 21.2 | 59.0 | 69.7 | 77.6 |

<sup>a</sup> Significance level for one-sided Wilcoxon signed-rank test: $p < 0.05$ (\*), $p < 0.01$ (\*\*), and $p < 0.001$ (\*\*\*)

**Table 3:** Pairwise test for significant improvement of the MS + RT score integration methods: *Our*, MetFrag 2.2 and Chain-graph. We show the p-values for testing the improvement of the row over the column method using a one-sided Wilcoxon signed-rank test. The test is performed over all top-*k* accuracy samples (datasets and ionization). MetFrag 2.2 and Chain-graph could not significantly outperform our framework. P-values ≥ 0.05 are marked with “n.s.”.

| Method<br>(MS + RT) | Top-1 |  | Top-20 |  |
| --- | --- | --- | --- | --- |
|  | Chain-graph | MetFrag 2.2 | Chain-graph | MetFrag 2.2 |
| *our* | $8.7 \cdot 10^{-24}$ | $6.1 \cdot 10^{-8}$ | $2.1 \cdot 10^{-13}$ | $1.5 \cdot 10^{-14}$ |
| Chain-graph | - | n.s. | - | $8.1 \cdot 10^{-04}$ |
| MetFrag 2.2 | $4.3 \cdot 10^{-08}$ | - | n.s. | - |

#### Influence of MS^2^ Scoring Method

Table 4 shows the performance using of our score integration framework for two difference MS^2^ scoring methods, MetFrag and IOKR (Sec. 3.3). Retention order information (MS + RT) can improve the identification performance in both cases, however the improvement with IOKR scores is lower.

**Table 4:** Top-*k* accuracies averaged across all datasets for two MS^2^-scorers.

| MS<sup>2</sup>-Scorers | Method | Top-1 | Top-5 | Top-10 | Top-20 |
| --- | --- | --- | --- | --- | --- |
| MetFrag | MS + RT | 21.3 | 52.9 | 64.0 | 74.3 |
|  | Only MS | 16.7 | 49.5 | 60.4 | 70.6 |
| IOKR | MS + RT | 26.7 | 52.1 | 62.5 | 70.3 |
|  | Only MS | 25.1 | 49.5 | 60.3 | 67.6 |

#### Influence of the Candidate Set

Here we study the effect of two commonly used ways of defining the candidate lists of molecular structures: First approach (“All”) includes all molecules with a matching exact mass to the list, and the second approach (“Correct MF”) only includes molecular structures matching the pre-determined molecular formula (e.g. SIRIUS (Dührkop *et al*., 2019) uses this approach). To determine the effect of the candidate set definition on our framework we modify the CASMI dataset, such that for a spectrum *i* only candidates are used that have the same molecular formulas as the correct structures. This leads to significantly reduced candidate sets: For the positive mode spectra the median number of candidates decreases from 919 to 238 and for the negative ones from 420 to 58. For the Massbank data we cannot do this modification, as the candidates are already restricted to structures with the correct molecular formula. Table 5 shows that the baseline performance using MetFrag MS^2^ scores (Only MS) improves after filtering of the candidates. Further improvement can be reached by using retention order information (MS + RT) even though the absolute improvement is slightly lower than without candidate filtering. For IOKR, we notice that RT information significantly improves the top-*k* accuracies when all matching exact mass candidates are used, whereas when the candidate sets only contain molecules with the correct, i.e. ground truth, molecular formula (MF), using RT information can only improve top-10 and top-20 accuracies.

**Table 5:** Top-*k* accuracies averaged on the CASMI data (pos. & neg.) using either MetFrag or IOKR as MS^2^-scorer for two different candidate sets: “All” molecules queried using a mass window; Only those with “correct molecular formula” (MF). The stars (*) represent the significant improvement over the Only MS (see Table 2 for details on the significance test).

| MS<sup>2</sup>-Scorer | Candidate Set | Method | Top-1 | Top-5 | Top-10 | Top-20 |
| --- | --- | --- | --- | --- | --- | --- |
| MetFrag | All | MS + RT | 14.6 (***) | 44.0 (***) | 54.6 (***) | 66.5 (***) |
|  |  | Only MS | 11.4 | 40.7 | 51.2 | 63.2 |
|  | Correct MF | MS + RT | 17.7 (***) | 48.4 (***) | 59.8 (***) | 71.0 (***) |
|  |  | Only MS | 13.1 | 46.0 | 56.9 | 68.7 |
| IOKR | All | MS + RT | 26.0 (***) | 48.0 (***) | 60.0 (***) | 69.1 (***) |
|  |  | Only MS | 24.4 | 46.0 | 58.4 | 65.5 |
|  | Correct MF | MS + RT | 30.6 | 52.3 | 66.2 (***) | 75.1 (*) |
|  |  | Only MS | 30.6 | 53.9 | 65.3 | 74.8 |

### 4.3 Missing MS^2^

In Figure 3 we show the identification accuracy using our score integration framework compared to the baseline (Only MS) when only some percentage of the MS features have an MS^2^ spectrum. The features without spectra only use the precursor mass as MS information (Sec. 3.6). We vary the percentage from 0% to 100% with 25%-point steps. The retention order weight *D* was optimized using the 100% setting. At 0% the score integration framework only uses the mass of the candidates and their predicted retention order for the ranking. In the absence of MS^2^ information, we observe a high performance gain for top-20. The more MS^2^ information we add, the smaller the gain in top-20 accuracy using the retention orders. The fact that RT is a weaker information than MS^2^ could explain this observation. The more MS^2^ are available, the less additional information RT can add. For the top-1 there is constant improvement for all MS^2^ %’s.

**Figure 3:**
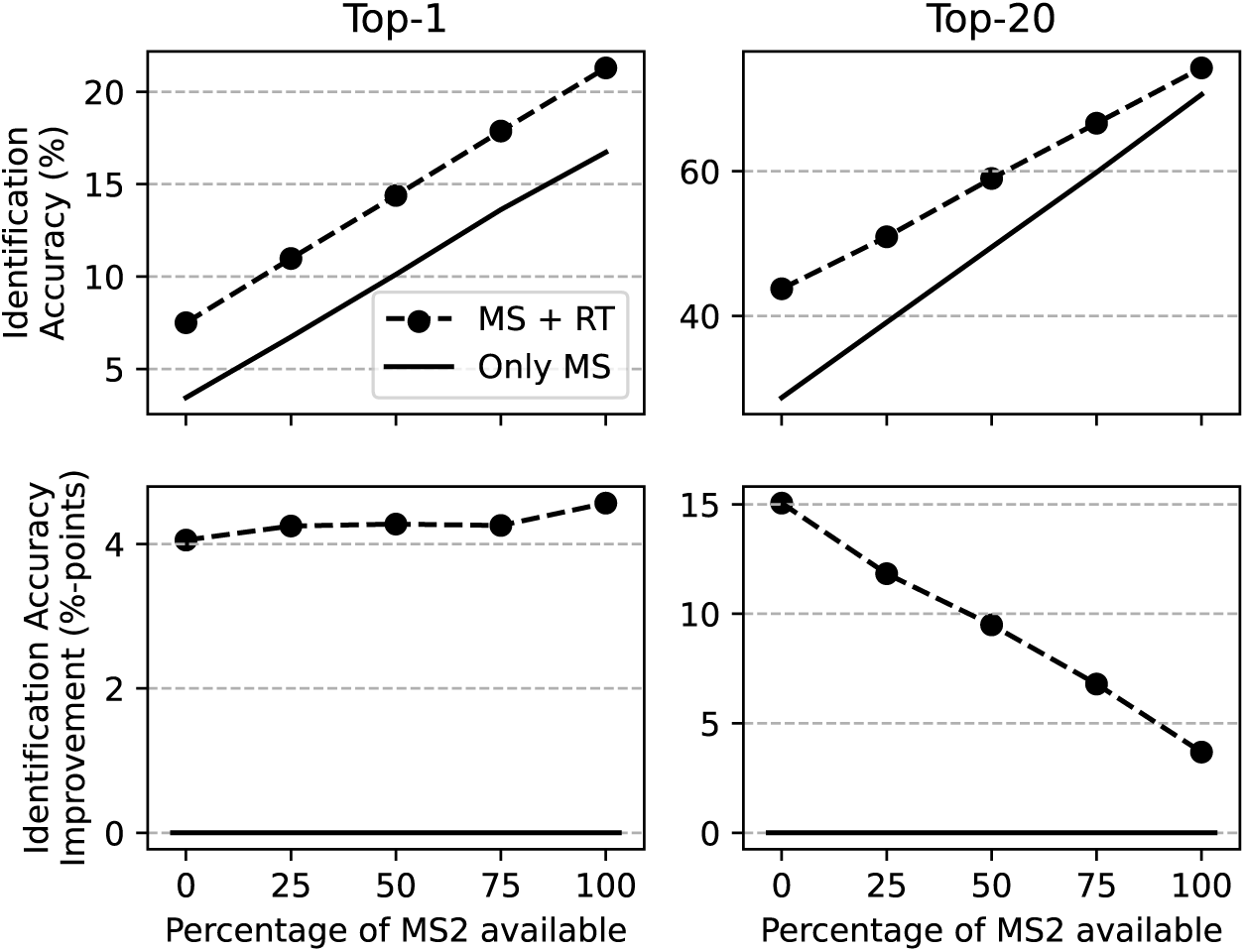
Top-*k* accuracies and improvements averaged across all datasets. Plots for different percentages of available MS^2^ spectra: 0% (only MS^1^ and RT) to 100% of MS^2^ spectra (previous experiments).

## 5 Discussion

In this paper we have put forward a rigorous probabilistic framework for the integration of mass spectrometry (MS) based candidate structure and retention order predictions. Our framework allows the use of any of the popular models, such as CSI:FingerID, IOKR or Metfrag for scoring candidate structures on MS data.

Out method takes into account the retention orders of all candidate structure pairs in distinct candidate lists through an approximated fully connected Markov random field model. It generally achieves higher quality structural annotations of observed MS features than using a single Markov chain as implied in the Bach *et al.* (2018) model. It also improves on the method of Ruttkies *et al.* (2016), which uses predicted retention times (RT), in three out of four experiments. For the latter approach, we believe using the RankSVM scores instead of the predicted LogP values could improve the performance. Both measures are proxies for retention behavior and our results show that the RankSVM predicts the retention order more accurately than the LogP values (see Table S2 in supplementary material). We also demonstrate, that our framework improves the identifications, if only a subset of the MS features come with an MS^2^ spectra. The framework is computationally efficient and can be trained using modest-sized datasets.

The amount of improvement using RT information was shown to depend on the dataset and MS^2^ scorer (here MetFrag or IOKR). This indicates that RT information rather fine tunes the ranking given by the MS^2^ scorer, for example by better tie-breaking. The underlying factors could be ambiguities in the candidate sets that can be only be resolved by RT or molecular features that cannot be predicted by MS. Stereo-chemistry is an obvious factor, but reliable annotations of stereo-chemistry are generally missing from RT databases and thus cannot currently be used for training better retention order prediction models. Thus improved modelling of stereo-chemistry features is an important open problem (Witting and Böcker, 2020). A further research direction could be to include the liquid chromatography (LC) system’s configuration, e.g. column or eluent, into the retention order prediction. As LC systems can be configured to separate certain molecular classes, this could provide additional information to certain molecular candidates. Also, using the LC peak shape to train a model directly predicting the retention order probabilities could be more accurate, e.g. by incorporating RT variance. However, such data is currently not part of the public RT databases.

## Acknowledgements

E.B. likes to thank Emma Schymanski for answering numerous questions on mass spectrometry and Sandor Szedmak for his helpful comments on the computational framework.

## Supplementary information

Supplementary data are available at *Bioarxiv* online.

## Footnotes

1 http://msbi.ipb-halle.de/~cruttkie/metfrag/MetFrag2.4.5-CL.jar

