## Supplementary Material for "Probabilistic Framework for Integration of Mass Spectrum and Retention Time Information in Small Molecule Identification"

### Contents

**S.1 Code and Data links** 1

**S.2 Hyper-parameter Estimation** 1

**S.3 Results** 1
S.3.1 Comparison of the Edge Potential Functions . . . . . . . . . . . . . . . . . . . . . . 1
S.3.2 Retention Order Prediction Model . . . . . . . . . . . . . . . . . . . . . . . . . . . . 2

**S.4 Input Kernels for the IOKR Models** 2

### S.1 Code and Data links

The code and data used in this publication are available at `https://github.com/aalto-ics-kepaco/msms_rt_score_integration`.

### S.2 Hyper-parameter Estimation

Algorithm 1 provides the pseudo-code for the hyper-parameter selection procedure, which is verbally described in Section 3.5, to determine the optimal retention order weight $D^*$ given a labeled training set $\mathcal{D}_{train}$. The algorithm can be easily extended to also determine the optimal parameter $k^*$ of a sigmoid function, by searching a grid of $(D, k) \in \mathbf{D} \times \mathbf{k}$ tuples. This latter extension was used to determine $(D^*, k^*)$ for the evaluation of the chain-graph approach (Sec. 3.6 and 4.2) using the Hinge-Sigmoid as edge-potential function.

### S.3 Results

This section contains additional results not shown in the main document.

#### S.3.1 Comparison of the Edge Potential Functions

In Table S1 we compare the metabolite identification performance of the different edge potential functions presented in Section 2.2.3. The results are discussed in Section 4.1.2. The sigmoid

**Algorithm 1**: Hyper-parameter Estimation Procedure. This procedure is applied to the sum- as well as max-marginals.

**Data**: $\mathbf{D}$ retention order weight ($D$) grid; $\mathcal{D}_{train}$ labeled set of MS-features with candidate set used to evaluate the performance of each grid value.

**Result**: $D^* \in \mathbf{D}$ retention order weight with the highest performance.

$s \leftarrow \{D : -1 \mid D \in \mathbf{D}\}$;

**for** $D \in \mathbf{D}$ **do**

  /* Get normalized marginals for tree sample $\mathbb{T}$ (Sec. 2.3.1 and 2.3.2). */

  **for** $t \in \{1, \ldots, L\}$ **do**

    $p(\cdot \mid T_t) \leftarrow$ `get_normalized_marginals`($\mathcal{D}_{train}$, $T_t$, $D$);

  /* Get average marginal (Sec. 2.3.1). */

  $\bar{p}(\cdot) \leftarrow$ `get_averaged_marginal`($\{p(\cdot \mid T_t)\}_{t=1}^{L}$);

  /* Evaluate the performance of $D$ on $\mathcal{D}_{train}$ (Sec. 3.5). */

  $s(D) \leftarrow$ `get_top20AUC_performance`($\bar{p}(\cdot)$, $\mathcal{D}_{train}$);

$D^* \leftarrow \arg\max_{D \in \mathbf{D}} s(D)$;

Table S1: Identification accuracies (top-k) for different edge potential functions. We use the max-marginal and $L = 128$ for the score integration. The accuracies are averaged across all datasets and ionizations. Both potential functions improve significantly ($p < 0.001$, one-sided Wilcoxon signed-rank test) over the *Only MS* setting.

| **Method** | **Edge-potential** | **Top-1** | **Top-5** | **Top-10** | **Top-20** |
|---|---|---|---|---|---|
| MS + RT | Sigmoid | 21.3 | 52.9 | 64.0 | 74.3 |
| | Step-function | 20.8 | 52.6 | 64.3 | 74.4 |
| Only MS | - | 16.7 | 49.5 | 60.4 | 70.6 |

function can outperform the step-function significantly for top-1 ($p < 0.001$) and top-5 ($p < 0.05$). As, on the other hand, the improvement of the step- over the sigmoid function is not significant, we decided to use the sigmoid function for the majority of our experiment.

#### S.3.2 Retention Order Prediction Model

Table S2 shows the retention order prediction accuracy of the RankSVM models for the different evaluation datasets (Sec. 3.1) evaluated on the random subsets used for our score integration experiments. It furthermore compares the RankSVM with the CDK XLogP in terms of their retention order modelling performance. The predicted LogP values are used by MetFrag 2.2 to predicted the candidates' retention times and subsequently re-rank them (Sec. 3.6). The RankSVM scores ($\mathbf{w}^T\phi_{ir}$, Sec. 2.2, *Edge Potential Function*) and the CDK XLogP values, can be considered as a proxy for retention order behavior.

### S.4 Input Kernels for the IOKR Models

This section contains a description of the $MS^2$ and fragmentation tree (FT) (Böcker and Dührkop, 2016) kernels used for the IOKR models described in Section 3.3 of the main document. FTs are a representation of the fragmentation process a molecule undergoes during the $MS^2$ analysis. The tree is deduced from the given $MS^2$ spectrum. Its nodes represent the predicted molecular formulas for each $MS^2$ peak. The edges are labeled with a predicted molecular formula of the loss between two peaks in the spectrum (Böcker and Rasche, 2008; Böcker and Dührkop, 2016). We

Table S2: The average pairwise prediction accuracy calculated for the correct molecular structures of each subsample. It expresses the agreement of the RankSVM and CDK XLogP with the observed retention orders.

| | | **Order prediction accuracy** | |
|---|---|---|---|
| **Dataset** | **Ionization** | RankSVM | CDK XLogP |
| CASMI 2016 | negative | 0.84 (± 0.04) | 0.79 (± 0.05) |
| | positive | 0.83 (± 0.04) | 0.72 (± 0.05) |
| EA (Massbank) | negative | 0.87 (± 0.02) | 0.81 (± 0.03) |
| | positive | 0.88 (± 0.02) | 0.75 (± 0.05) |

used 16 kernels for our models of which one, PPK, is a spectra kernel and 15 are FT kernels. An overview of the kernels can be found in Table S3.

Table S3: Description of the $MS^2$ and fragmentation tree kernels used for the IOKR models. Read (Dührkop, 2018; Dührkop *et al.*, 2015) for further details. Abbreviations: Molecular formula (MF), Fragmentation tree (FT). The nodes, in a FT, are associated with spectra peaks and the edges are associated with losses.

| **Abbreviation** | **Name** | **Description** |
|---|---|---|
| CPJXB | Common Path Joined Binary | Number of paths with equal union of losses |
| CPJ | Common Path Joined | Count of length two paths with the same loss |
| LC | Loss count | Count of each loss in the FT |
| LI | Loss Intensity | Intensity weighted counts of common losses |
| LPC | Loss Pair Count | Count for each pair of consecutive losses in the FT |
| MLIP | Maximum Loss in Path | Maximum frequency if each loss in any path of the FT |
| NB | Node Binary | Number of nodes sharing the same molecular formula |
| NI | Node Intensity | Intensity weighted variant of NB |
| NSFLC2 | Node Loss Interaction | Counts the common paths and weights them by comparing the MF of their terminal fragments |
| RLB | Root Loss Binary | Number of common root-losses |
| RLI | Root Loss Intensity | Intensity weighted variant of RLB |
| UFS1 | Substructure in Losses and Leafs | Number of times a predefined set of MFs is preserved in a path or cleaved of intact |
| UFS3 | - | Same as UFS1 but values taken to the power of three |
| WFPC | Weighted Fingerprint Path | Count paths in the FT that correspond to certain molecular properties |
| WNSF | Weighted Substructure Counting | Count set molecular substructures present in the training and weight by their occurrence |
| PPKr | Probability Product Kernel | Probability product kernel computed on the peaks of preprocessed $MS^2$ spectra |
